# A beary good genome: Haplotype-resolved, chromosome-level assembly of the brown bear (*Ursus arctos*)

**DOI:** 10.1101/2022.06.17.496447

**Authors:** Ellie E. Armstrong, Blair W. Perry, Yongqing Huang, Kiran V. Garimella, Heiko T. Jansen, Charles T. Robbins, Nathan R. Tucker, Joanna L. Kelley

## Abstract

The brown bear (*Ursus arctos*) is the second largest and most widespread extant terrestrial carnivore on Earth and has recently emerged as a medical model for human metabolic diseases. Here, we report a fully-phased chromosome-level assembly of a male North American brown bear built by combining Pacific Biosciences (PacBio) HiFi data and publicly available Hi-C data. The final genome size is 2.47 Gigabases (Gb) with a scaffold and contig N50 length of 70.08 and 43.94 Mb, respectively. BUSCO analysis revealed that 94.5% of single-copy orthologs from mammalia were present in the genome (the highest of any ursid genome to date). Repetitive elements accounted for 44.48% of the genome and a total of 20,480 protein coding genes were identified. Based on whole genome alignment, the brown bear is highly syntenic with the polar bear, and our phylogenetic analysis of 7,246 single-copy BUSCOs supports the currently proposed species tree for Ursidae. This highly contiguous genome assembly will support future research on both the evolutionary history of the bear family and the physiological mechanisms behind hibernation, the latter of which has broad medical implications.

**Significance:** Brown bears (*Ursus arctos*) are the most widespread, large terrestrial carnivore on the planet and represent an interesting example of speciation through hybridization, as well as a medical model for sedentary lifestyle-related disease. Although a previous genome for a brown bear has been published, the reported contig N50 was low (only ∼530 kb), despite being scaffolded into putative chromosomes. Genomes of this quality limit the accuracy of analyses which rely on long contiguous stretches of the genome to be assembled (such as with many demographic analyses) as well as attempts at connecting genotype to phenotype (such as in association analyses). In order to support studies on both the complex hybridization history of the brown bear and investigations into medically-relevant phenotypes, we generated a fully-phased, chromosome-level assembly from a male grizzly bear. The genome has a total size of 2.47 Gb and 90% of the genome is contained in 36 scaffolds, roughly corresponding to one autosome per scaffold. This high-quality genome will enable studies across a variety of disciplines, including conservation, evolution, and medicine.

## Introduction

Brown bears (*Ursus arctos*) are a historically wide-ranging species, formerly occupying habitat from the southern tip of North America, across most of Asia and Europe, and along the northernmost tip of Africa (Iucn & IUCN 2016). However, as the second largest extant terrestrial carnivore, brown bears have seen extensive reductions in their range and even total extirpations in some regions due to habitat loss, climate change, and human-wildlife conflict (Albrecht et al. 2017). As top predators, brown bears also play an important role in ecosystem function (Duffy 2003). Brown bears are interesting ecological models that show local adaptations in both diet (Bojarska & Selva 2012), morphology (Colangelo et al. 2012; Sato et al. 2011), and other life history traits (Ferguson & McLoughlin 2000).

Brown bears have emerged as a model species for population genomics and speciation due to their interesting (and not fully resolved) demographic history, which contains signals of both incomplete lineage sorting and post-speciation hybridization (Barlow et al. 2018; Cahill et al. 2013, 2015; Kumar et al. 2017; Miller et al. 2012). Additionally, brown bear hibernation has been proposed as a medical model for several diseases, including diabetes and insulin resistance (Rigano et al. 2017) and conditions related to sedentary-lifestyles (Fröbert et al. 2020). While several genome assemblies for the brown bear have been published to date (Taylor et al. 2018), these assemblies have low contiguity (i.e., contig N50 = ∼530 kilobases (kb)) which limits their value when being used to study brown bear biology. In order to improve the power and breadth of future research for the brown bear, we present a fully-phased, chromosome-level assembly from a male brown bear built with Pacific Biosciences (PacBio) HiFi data and scaffolded with publicly available Hi-C data. We analyze this genome for quality and completeness, report on improved annotation statistics, and compare it with other publicly available bear genomes for diversity, demographic history, and repetitive element composition.

## Results and Discussion

### Genome quality and continuity

Utilizing the trio-binning method via Hifiasm, we generated a phased assembly with one haplotype phase (Hifiasm-Hap1) totaling 2.47 Gigabases (Gb) and the other (Hifiasm-Hap2) totaling 2.46 Gb (Table S1) with a contig N50 of 48.3 and 48.2 Mb, respectively. The contig L90 indicated that 55 and 54 contigs made up 90% of the total genome. Note that for long read assemblies that do not incorporate a scaffolding step, only contig statistics are reported since no scaffolds are built. After incorporation of Hi-C data, the scaffold and contig N50 for hap1 (Hifiasm-Hap1 + HiC) and hap2 (Hifiasm-Hap1 + HiC) were 70.5 and 45.6 Mb, and 70.1 and 43.9 Mb, respectively. The slight decrease in contig N50 after Hi-C data incorporation likely indicates that some misassemblies were present in the original PacBio HiFi assembly. The final composite assembly includes the autosomes and unplaced scaffolds from Hifiasm-Hap2 + HiC, the putative Y chromosome scaffolds from Hifiasm-Hap2, and the major X scaffold from Hifiasm + HiC Hap1. The composite assembly had a scaffold and contig N50 of 70.1 and 43.9 Mb and a scaffold L90 of 36 (Table S1), indicating that 90% of the assembly is contained in 36 scaffolds. Given the statistics of the final assembly, it is likely that most autosomes and the X chromosome are contained in approximately one scaffold, since the diploid Ursine karyotype is 37 (Nash et al. 1998; Wurster-Hill & Bush).

Previously, approximately 1.9 Mb of putative Y chromosome scaffolds were identified in a previous version of the polar bear assembly (Bidon et al. 2015), while the most recent polar bear assembly contains approximately 1.6 Mb of putative Y scaffolds (see GCF_017311325.1 assembly report via UCSC browser). After removing a misassembly, we identified a total of approximately 9.9 Mb of putative Y scaffolds based on alignment to the Y scaffolds in the polar bear assembly (Table S2). However, it is likely that there are still some mis-assemblies within this region due to the repetitive nature of mammalian Y chromosomes (Li et al. 2013).

The final composite assembly also improved upon the two previously published assemblies for *Ursus arctos*, which had a scaffold/contig N50 of 36.7/0.5 Mb and 72.2/0.5 Mb, respectively (Table S3). Although the assembly produced by DNAZoo has a slightly better scaffold L90 (32; Table S3), the contig N50 is improved in our assemblies by approximately 88x. Undoubtedly, Hi-C data from a male bear (current Hi-C data is from a female) and/or additional long read data from a male bear will provide further improvements and resolution to this assembly.

BUSCO analyses revealed that each bear haplotype phase and the composite assembly had from 96.3-96.5% of expected complete genes (Table S4). We observed no changes in BUSCO scores when incorporating Hi-C data (Table S4), revealing that any joins or misassemblies did not impact these genic regions. The BUSCO scores from the assemblies produced here are the highest scores across any currently published bear assembly (Table S5), further indicating that the final assembled genomes are of high quality.

### Genomic synteny

In order to investigate the synteny between our genome and the polar bear (the closest relative to the brown bear with an estimated divergence date of more than one million years (Lan et al. 2022; Bidon et al. 2014; Cronin et al.)), we performed a whole-genome alignment. Both the polar bear and the brown bear are ursine bear species, which are known to have a stable karyotype of 2n = 74 (Nash et al. 1998; Wurster-Hill & Bush). This is in agreement with the alignment produced here for the polar and brown bear (Figure 1a), which showed no major chromosomal rearrangements. For both the polar and brown bear, most of all 36 autosomes and the X chromosome appear to be represented primarily by one scaffold each, reiterating the quality of the assembly (Table S1).

**Figure 1:**
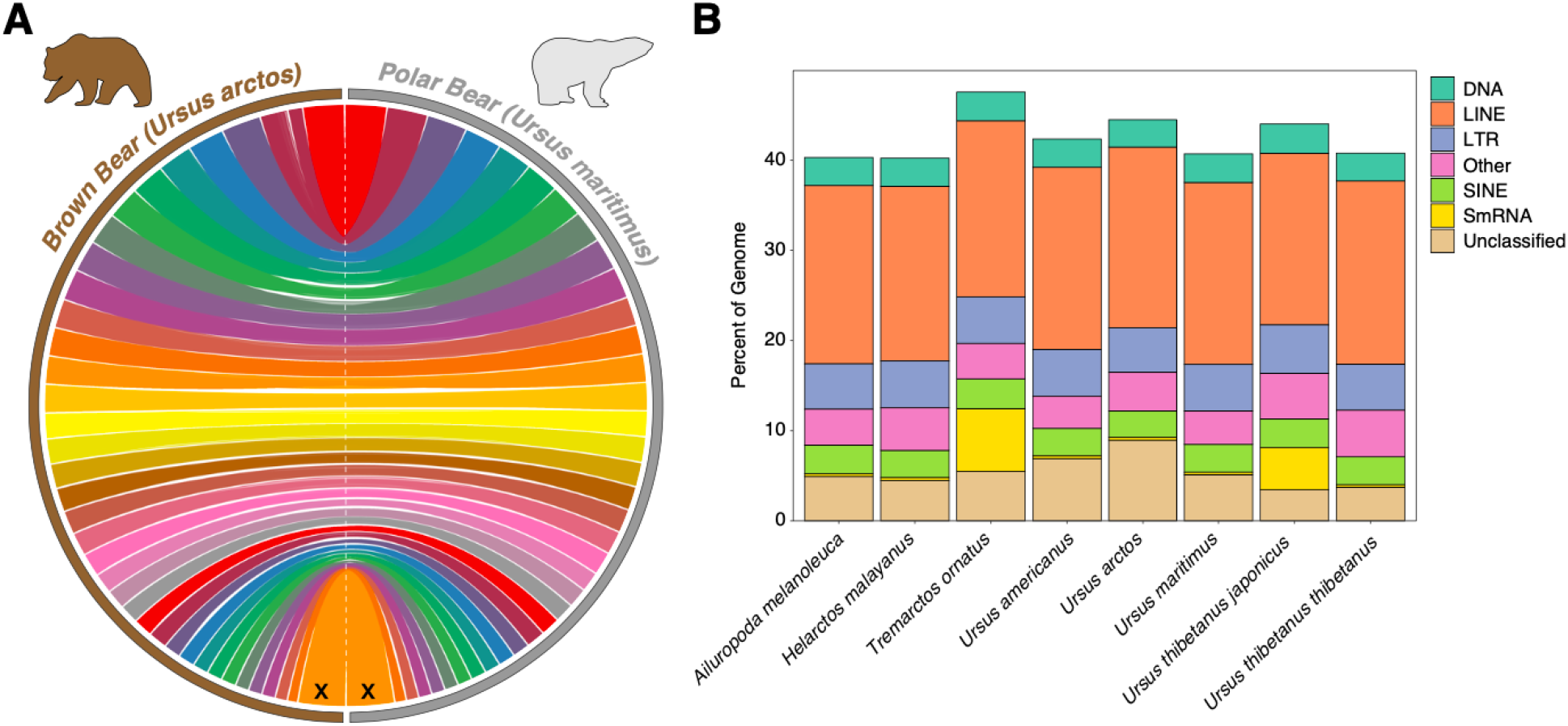
A) Whole genome alignment between the brown bear and the polar bear containing all predicted autosomal scaffolds and the X chromosome. B) Repeat content across Ursidae.

### Repetitive content

Across all bears, a total of 40.21-47.54% of the genome was made up of repetitive elements (Figure 1b). Across most repetitive element classes, a majority of the bears had comparable numbers, but differed most substantially in the total amount of ‘Small RNA’, ‘Unclassified’, and ‘Other’ (comprising satellites, simple repeats, and low complexity regions) (Figure 1b). Consistent with previous results, we found that long interspersed nuclear elements (LINEs) represented the largest percentage of repetitive elements in the Ursidae family (Zhu et al. 2020; Srivastava et al. 2019), however, one study reported fewer total repeats in the American black bear (*Ursus americanus*) and a greater number of repeats overall in the giant panda (*Ailuropoda melanoleuca*) (Srivastava et al. 2019). Interestingly, while most Ursid species contain a relatively low percent of small RNA repetitive elements, we find expansions of this repetitive element class in the Andean bear (*Tremarctos ornatus*) and the Japanese black bear (*Ursus thibetanus japonicus*) (Figure 1b). As these genomes have some of the lowest quality scores (see Table S3 and Table S4) improved assemblies will be needed to investigate whether this is an artifact of misassembly or reflects the actual repetitive element content.

### Gene content

A total of 29,516 genes and pseudogenes were predicted in the genome assembly by the National Center for Biotechnology Information’s annotation pipeline (*https://www.ncbi.nlm.nih.gov/genome/annotation_euk/Ursus_arctos/102/*). Of these, a total of 20,480 were protein coding, 5,160 were non-coding, 3,630 were pseudogenes, and 219 were immunoglobulin gene segments. Compared to the previously annotated genome for *Ursus arctos* (Taylor et al. 2018), this assembly adds 632 protein coding genes, reduces the number of non-coding and pseudogene sequences by 1,901 and 41, respectively, and improves the number of immunoglobulin gene segments by 100. This evidence, along with evidence provided by BUSCO (see section: *Quality and continuity*), indicates that the more complete and contiguous assembly has resulted in a more accurate assembly and annotation of gene regions.

### Phylogenetics

Using a total of 7,246 single-copy orthologs from the mammalia_odb10 BUSCO dataset, we generated a consensus tree for Ursidae (Figure 2a). Previous studies have found a substantial amount of gene tree-species tree discordance due to possible incomplete lineage sorting (ILS) and/or post-speciation hybridization (Barlow et al. 2018; Cahill et al. 2013, 2015; Kumar et al. 2017). While most of these studies have focused on the potential hybridization history of the brown bear and polar bear, recent work has suggested that ILS and post-speciation gene flow may be more prominent throughout the clade than previously expected (Kumar et al. 2017). Our results reiterate the basic species-tree topology and quartet scores revealed a high amount of gene tree-species tree discordance across the Ursid topology (Figure 2A). Reiterating the results from Kumar et al. 2017, we found strong signals of discordance both among American black bears, brown bears and polar bears (45-54% of genes supported the main tree topology in this lineage), but also in the Asiatic bear lineages (40-50%). However, while previous studies have relied on consensus sequence generation, *de novo* assemblies and reference-free alignments like those performed here avoid mapping and consensus generation-related errors when performing evolutionary inference (Prasad et al. 2022; Gopalakrishnan et al. 2017; Armstrong et al. 2020; Westbury et al. 2021). Utilization of such methods may be essential for understanding evolution in young lineages like the bears, with high degrees of ILS and hybridization.

**Figure 2:**
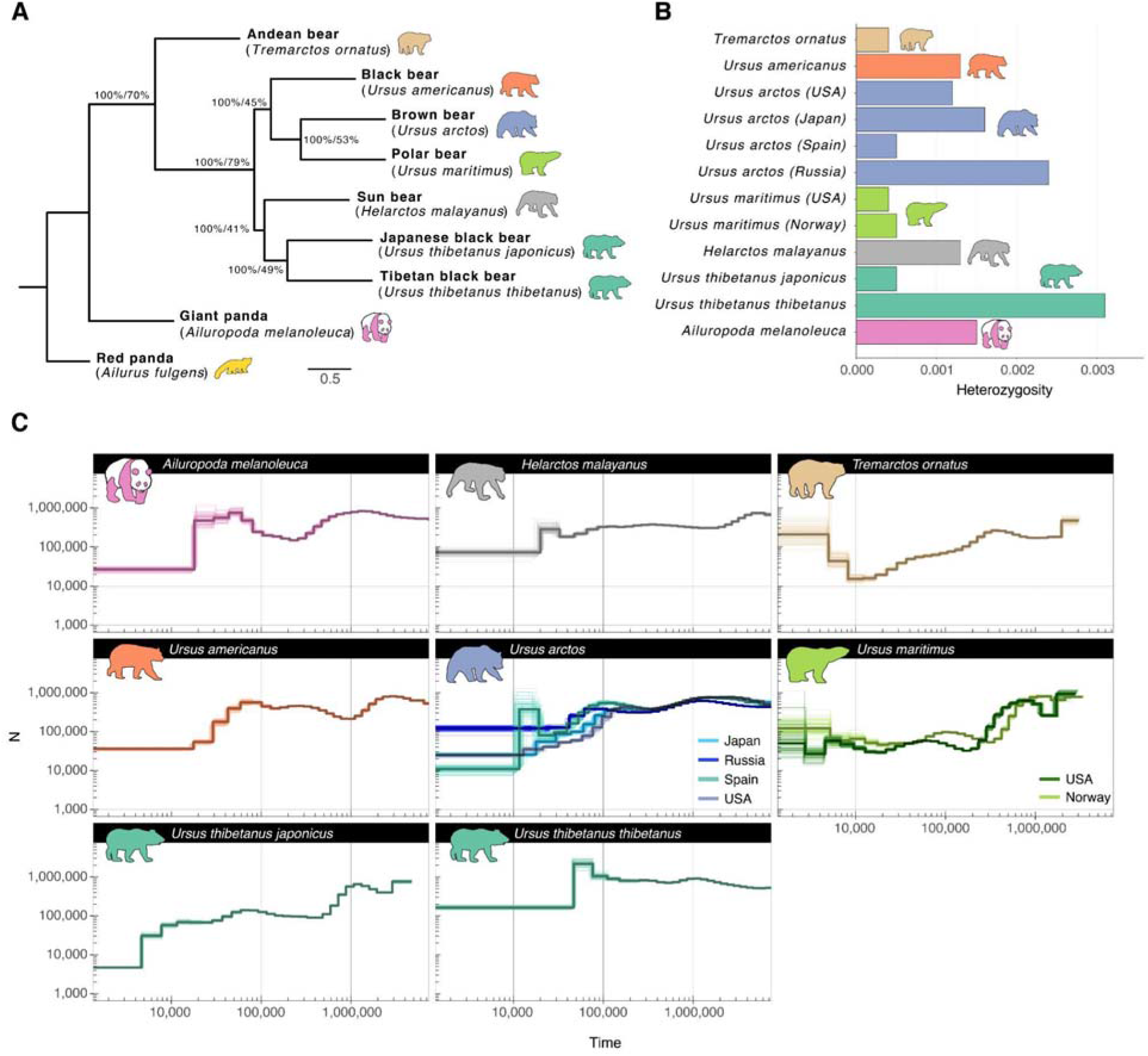
A) Consensus phylogenetic tree generated from 7,246 single-copy orthologs. B) Observed heterozygosity across various bear species and subspecies as calculated by angsd. C) PSMC estimates of effective population size over time for focal bear species and populations. Thick bold lines indicate estimates from the full dataset, while thinner lines indicate 100 individual bootstrap replicates.

### Heterozygosity

In order to investigate the relative diversity across bear species, we estimated observed heterozygosity for all species of bear for which whole-genome sequence data and assemblies are available. We also included various subspecies and/or populations (see Table S6 for details). We found that even within bear species, heterozygosity values vary widely depending on population. Similar to previous results, we found that the Apennine brown bears from Spain had the lowest heterozygosity values ((Endo et al. 2021); Figure 2b). The other brown bear tested from Europe had higher values than those from the isolated Hokkaido brown bear (ssp *lasiotus*) population. Polar bears, the Japanese black bear, and the sun bear also show remarkably low heterozygosity values, similar to those from the endangered Apennine brown bears. The Tibetan black bear (*Ursus thibetanus thibetanus*) showed the highest heterozygosity across any of the other bear species, especially compared to its island counterpart, the Japanese black bear. However, there is limited information on the connectivity and history of the mainland Asiatic black bear across its range, so additional individuals should be sequenced to establish if this estimate is representative of the larger population/species.

### Demographic history

We investigated the demographic history across the bear clade using Pairwise Sequentially Markovian Coalescent (PSMC). Our analyses are mostly consistent with previous investigations of demographic history in bears, including an increase in *N*_*e*_ approximately 120 kya for most mainland continental species (Figure 2C). Interestingly, although previous analyses have shown this *N*_*e*_ increase to be apparent in the Alaskan populations of the North American brown bear (Endo et al. 2021; Miller et al. 2012), we do not observe this in bears sampled from the lower 48 states. Moreover, inconsistencies with previous estimates of total *N*_*e*_ across species appear to be attributable to differences in the selected mutation rate. Here, we predict a nearly doubled *N*_*e*_ compared to previous results (Cahill et al. 2013; Kumar et al. 2017; Endo et al. 2021; Zhu et al. 2020; Liu et al. 2014; Lan et al. 2022), which is consistent with expectations for having a mutation rate that is approximately halved (Nadachowska-Brzyska et al. 2016). This mutation rate is likely more accurate since it was calculated directly through trio sequencing (Wang et al. 2022).

## Materials and Methods

### Sample acquisition and library preparation

In order to build a phased genome assembly using a trio-binning strategy, we collected blood samples from a bear trio (Adak, offspring; Oakley, mother; John, father). Protocols were followed according to (Joyce-Zuniga et al. 2016). All procedures were approved by the Washington State University Institutional Animal Care and Use Committee (IACUC) under protocol number ASAF 6546.

For short read whole genome sequencing of parental DNA, genomic DNA was isolated from frozen blood using the Gentra Puregene kit (Qiagen). PCR-free WGS libraries prepared at the Genomics Platform of the Broad Institute and were paired-end sequenced (2×150 bp) on a HiseqX to an estimated depth of 30X.

For long-read WGS, high molecular weight DNA was extracted from 3mL of fresh blood. Before DNA extraction, red blood cells were lysed using the RBC lysis buffer from the Gentra Puregene kit (Qiagen), and white blood cells were pelleted and washed. DNA from these cells was isolated using the Monarch HMW DNA Extraction Kit for Tissue (New England Biosciences, T3060). For PacBio library preparation, ≥3 ug of high molecular weight genomic DNA was sheared to ∼15 kb using the Megaruptor 3 (Diagenode B06010003), with DNA repair and ligation of PacBio adapters accomplished with the PacBio SMRTbell Express Template Prep Kit 2.0 (100-938-900). Incomplete ligation products were removed with the SMRTbell Enzyme Clean Up Kit 2.0 (PacBio 101-938-500). Libraries were then size-selected for 15 kb +/- 20% using the PippinHT with 0.75% agarose cassettes (Sage Science). Following Qubit dsDNA High Sensitivity assay quantification (Thermo Q32854), libraries were diluted to 60 pM per SMRT cell, hybridized with PacBio V5 sequencing primer, and bound with SMRT seq polymerase using Sequel II Binding Kit 2.2 (PacBio 101-908-100). CCS sequencing was performed on the Sequel IIe using 8M SMRT Cells (101-389-001) and the Sequel II Sequencing 2.0 Kit (101-820-200), PacBio’s adaptive loading feature was used with a 2 hour pre-extension time and 30 hour movie time per SMRT cell. Initial quality filtering, basecalling, adapter marking, and CCS error correction was done automatically on board the Sequel IIe. Sequencing yielded an estimated depth of coverage of 32X.

### Genome assembly

The haplotype-resolved assemblies were built using Hifiasm ((Cheng et al. 2021), and yak (https://github.com/lh3/yak/releases/tag/v0.1) following the documentation (see https://hifiasm.readthedocs.io/en/latest/trio-assembly.html#trio-binning-assembly). Briefly, yak is used for collecting parent-specific *k*-mer distributions with parental short reads. These *k*-mer distributions are then used for binning the (CCS) long reads of the offspring into paternal-specific and maternal-specific reads. Hifiasm is then used with the appropriately partitioned reads for constructing the haplotype-specific assemblies (paternal and maternal). The pipeline (https://github.com/broadinstitute/long-read-pipelines/blob/3.0.39/wdl/tasks/Hifiasm.wdl) performing this Hifiasm step is written in WDL.

After the trio-phased assembly was built using Hifiasm (Cheng et al. 2021), we subsequently used publicly available Hi-C data for the brown bear (courtesy of DNAZoo: DNAZoo.org) to further scaffold the assembly. To incorporate this data, we used Juicer (Durand et al. 2016) according to the standard DNA Genome Assembly Cookbook instructions (https://aidenlab.org/assembly/manual_180322.pdf). We used both haplotypes generated by Hifiasm in the previous step as input (separately). We then used the 3D-DNA pipeline (Dudchenko et al. 2017) to generate a draft assembly for both haplotypes generated from Hifiasm.

In order to identify putative sex chromosomes in each haplotype we used BLAST (Altschul et al. 1990) to identify which scaffolds/contigs in our genomes best aligned to the polar bear Y scaffolds (ASM1731132v1). The X scaffold was identified using whole genome alignments (see *Whole genome alignment* below). We found evidence that the male bear haplotype (Hifiasm-Hap2 + HiC) contained a misassembly of the Y and X chromosomes. In order to correct for this, we removed the two scaffolds containing BLAST hits from the Y chromosome and reincorporated the raw components of these scaffolds from the Hifiasm-Hap2 assembly into this genome. Last, in order to make a mappable genome that contained both sex chromosomes, we identified the major X chromosome scaffold from Hifiasm-Hap1 + HiC and incorporated it into the assembly. For our purposes, we refer to this as the ‘composite’ assembly.

### Quality and continuity assessment

We assessed the continuity and quality of each genome first using the Assemblathon2 scripts (Bradnam et al. 2013) followed by Benchmarking Universal Single-Copy Orthologs (BUSCOv5.3.0; (Simão et al. 2015) analysis. We analyzed all available bear assemblies using the *mammalia_odb10* datasets, with flags ‘--augustus’ and ‘-m genome’. For each species/subspecies, we selected the assembly with the best statistics from the Assemblathon2 and BUSCOv5.3.0 results to be used in the phylogenetics, repeat content analysis, and demographic history analyses. To see a complete description of which genomes were used, please refer to Table S6.

### Repeat content

In order to assess the relative repeat content across the bear genomes, we used a combination of homology-based repeat finding, as well as *de novo* repeat finding. Briefly, we first used RepeatMaskerv4.0.9 (Smit et al. 1996) to mask repeats based on known repeat databases using flags ‘-species Ursidae’, ‘-a’, and ‘-gccalc’ (Smit et al. 1996; Jurka et al. 2005). We then used the partially masked genome generated in the previous step as input to RepeatModeler v1.0.11 BuildDatabase, and subsequently performed *de novo* repeat finding using RepeatModelerv1.0.11 (Smit & Hubley 2008). Last, a masked file with both known and de novo repeats was produced by running RepeatMasker v4.0.9 with the flags ‘-gccalc,’ and ‘-a’, and a final library produced from the previous step as input with the initial masked file. Total repeat content was calculated by adding the values from the initial and de novo steps. Repeat content was visualized and plotted in R (Team & Others 2013) using ggplot2v.3.3.6 (Wickham 2011).

### Whole genome alignment

The brown bear genome assembly (composite) was aligned to the polar bear genome assembly (ASM1731132v1) in order to investigate assembly completeness, as well as genomic synteny. Briefly, genomes were aligned following scripts from https://github.com/mcfrith/last-genome-alignments using LASTv921 (Kiełbasa et al. 2011). Genome alignment was visualized using the CIRCA software (http://omgenomics.com/circa) by plotting only the major scaffold aligning to the putative 36 autosomes in the polar bear and the major X chromosome scaffold (see GCF_017311325.1 assembly report via UCSC browser). The major alignment was determined as the scaffold belonging to the query assembly (brown bear) that comprised a majority of the alignments to the putative polar bear chromosomes.

### Phylogenetics

We built a phylogenetic tree for all the members of Ursidae that have a genome assembly using the single-copy BUSCOs (see *Quality and continuity assessment*). We first extracted all single-copy BUSCOs generated with the mammalia_odb10 dataset, since this dataset resulted in higher numbers of complete, single-copy BUSCO’s across all *de novo* bear assemblies. Only genes which had a representative sequence from each species/subspecies were included. Each gene was then aligned using MAFFTv.7.490 (Katoh & Standley 2013) with the flags ‘--ep 0’, ‘--genafpair’, and ‘--maxiterate 1000’. Alignments were then trimmed using Gblocks v.091b (Castresana 2000) with flag ‘-t D’. Resulting files were then used as input into IQ-TREE 2 v. 2.1.3 (Minh et al. 2020) with flags ‘-bb 1000’, ‘-nt AUTO’, and ‘-m GTR+I+G’. Lastly, we concatenated the maximum likelihood trees and built a species tree using ASTRAL-III v5.7.8 (Mirarab et al. 2014; Zhang et al. 2018) with flags ‘-gene-only’ and ‘-t 2’ to annotate the tree. The resulting tree was then plotted in FigTree v1.4.3 (Rambaut 2007) and the tree manually rooted on *Ailurus fulgens* (red panda).

### Demographic history

We used the pairwise sequentially Markovian coalescent method to investigate demographic history across Ursidae, (Li & Durbin 2011). We analyzed data representing species, subspecies, and distinct populations of bears for which whole-genome sequencing data was available (see Table S6). Briefly, each genome was indexed using BWA *index* with flags ‘-a bwtsw’, and short-read data subsequently mapped using BWA-MEM. SAM files were converted to BAM format and sorted and an index generated. Subsequently, variant sites were called according to the suggested commands (see https://github.com/lh3/psmc). We used a minimum depth of 10 and a maximum depth of 100 for all samples except for the polar bear from Alaska, which was run with a minimum depth of 5 and a maximum depth of 50 due to it having a lower average sequencing depth (see Table S6).

We next generated PSMC curves with 100 bootstraps using the suggested parameters linked above, with a mutation rate of 0.9225 × 10-9 per bp per year (Wang et al. 2022) and a generation time of 10 years. Although there are a number of different generation times used for bears, we selected a generation time of 10 because we believe this to be a conservative estimate of generation time based on previous field studies (McLellan et al. 2017). We do note however, that small shifts in generation time are unlikely to impact the results of PSMC and only doubling this time will considerably impact results (Nadachowska-Brzyska et al. 2016). PSMC results were imported into R using psmcr v. 0.1-4 (see github.com/emmanuelparadis/psmcr) and plotted using ggplot2 v.3.3.6 (Wickham 2011).

### Heterozygosity

We estimated heterozygosity for each unique subspecies of bear (for individuals see *Demographic history*). Using the previously generated bam files as input in the program angsd v.0.931 (Korneliussen et al. 2014), we set the reference and the ancestral sequence as the genome assembly for each respective species, along with the flags ‘-GL 1’, ‘-dosaf 1’, ‘-fold 1’, ‘-minQ 20’, and ‘-minmapq30’. We generated folded spectra using the reference as the ancestral sequence, since the ancestral sequence is unknown. Subsequently, we ran the command realSFS within angsd and subsequently calculated the heterozygosity in R (Team & Others 2013).

## Data Availability

All data associated with this project has been deposited under NCBI Bioproject Accession PRJNA807323. Intermediate assemblies and phased haplotypes available at [DOI TBD].

## Acknowledgements

This research used resources from the Center for Institutional Research Computing at Washington State University. This work was supported by an NSF Office of Polar Programs (OPP) grant [award number 1906015] to JLK, an NSF OPP Post-doctoral Fellowship [award number 2138649] to BWP, the International Association for Bear Research and Management, Interagency Grizzly Bear Committee, USDA National Institute of Food and Agriculture (McIntire-Stennis project 1018967), Mazuri Exotic Animal Nutrition, and the Raili Korkka Brown Bear Endowment, Nutritional Ecology Endowment, and Bear Research and Conservation Endowment at Washington State University. EEA is a Washington Research Foundation Postdoctoral Fellow. NRT is supported by 5K01HL140187 from the National Institutes of Health. We also thank the volunteers and staff of the Washington State University Bear Center.

## Supplementary Tables & Figures

**Table S1:** Assembly statistics for intermediate and final brown bear assembly produced in this study. The N50 is the contig or scaffold length at which 50% of the genome is covered. The L90 is the number of contigs or scaffolds that comprise 90% of the genome length. Both statistics assume scaffolds/contigs are ordered from largest to smallest.

|  | **Total**<br>**Assembly**<br>**Length** | **Number**<br>**of**<br>**Contigs** | **Contig**<br>**N50** | **Contig**<br>**L90** | **Number**<br>**of**<br>**Scaffolds** | **Scaffol**<br>**d N50** | **Scaff**<br>**old**<br>**L90** |
| --- | --- | --- | --- | --- | --- | --- | --- |
| **Hifiasm-**<br>**Hap1** | 2,470,812,077 | 657 | 48,306,755 | 55 | N/A | N/A | N/A |
| **Hifiasm-**<br>**Hap2** | 2,459,149,093 | 534 | 48,190,235 | 54 | N/A | N/A | N/A |
| **Hifiasm-**<br>**Hap1 +**<br>**HiC** | 2,470,895,077 | 1,089 | 45,557,465 | 69 | 923 | 70,474,<br>880 | 35 |
| **Hifiasm-**<br>**Hap2 +**<br>**HiC** | 2,459,274,093 | 1,145 | 43,943,000 | 79 | 895 | 70,076,<br>652 | 36 |
| **Composite** | 2,474,241,652 | 1,116 | 43,943,000 | 77 | 901 | 70,076,<br>652 | 36 |

**Table S2:** Top BLAST hits of polar bear Y scaffolds against male haplotypes from this study. Scaffolds in the left column are scaffolds that were incorporated into composite assembly.

| **Polar bear scaffold** | **Hifiasm-Hap2 + HiC top hit** | **Score (Bits) / E value** | **Hifiasm-Hap2 top hit** | **Score (Bits) / E value** |
| --- | --- | --- | --- | --- |
| NW_024423324 | HiC_scaffold_1 | 47125/0.0 | h1tg000062l | 47125/0.0 |
| NW_024423325 | HiC_scaffold_2 | 31504/0.0 | h1tg0000139l | 31504/0.0 |
| NW_024423326 | HiC_scaffold_2 | 15459/0.0 | h1tg000081l | 15459/0.0 |
| NW_024423327 | HiC_scaffold_705 | 16325/0.0 | h1tg000062l,<br>h1tg000081l | 16325/0.0<br>16271/0.0 |
| NW_024423328 | HiC_scaffold_2 | 8100/0.0 | h1tg0000139l | 8100/0.0 |
| NW_024423329 | HiC_scaffold_1 | 10595/0.0 | h1tg000062l | 10595/0.0 |
| NW_024423330 | HiC_scaffold_1 | 7675/0.0 | h1tg000062l | 7675/0.0 |
| NW_024423331 | HiC_scaffold_1 | 8250/0.0 | h1tg000062l | 8250/0.0 |
| NW_024423332 | HiC_scaffold_2 | 6806/0.0 | h1tg0000101l | 9777/0.0 |
| NW_024423333 | HiC_scaffold_2 | 12198/0.0 | h1tg000081l | 12198/0.0 |
| NW_024423334 | HiC_scaffold_2 | 1491/0.0 | h1tg000081l,<br>h1tg000062l | 1491/0.0<br>1485/0.0 |
| NW_024423335 | HiC_scaffold_1 | 44754/0.0 | h1tg000062l | 44754/0.0 |
| NW_024423336 | HiC_scaffold_1 | 33972/0.0 | h1tg000062l | 33972/0.0 |
| NW_024423337 | HiC_scaffold_1 | 27235/0.0 | h1tg000062l | 27235/0.0 |
| NW_024423338 | HiC_scaffold_2 | 26596/0.0 | h1tg000081l | 26596/0.0 |
| NW_024423339 | HiC_scaffold_1 | 17494/0.0 | h1tg000062l | 17494/0.0 |
| NW_024423340 | HiC_scaffold_658 | 79106/0.0 | h1tg000081l,<br>h1tg000062l | 79106/0.0<br>54492/0.0 |
| NW_024423341 | HiC_scaffold_2 | 16397/0.0 | h1tg000081l | 16397/0.0 |
| NW_024423342 | HiC_scaffold_2 | 21943/0.0 | h1tg000062l,<br>h1tg000081l | 22018/0.0<br>21943/0.0 |

**Table S3:**
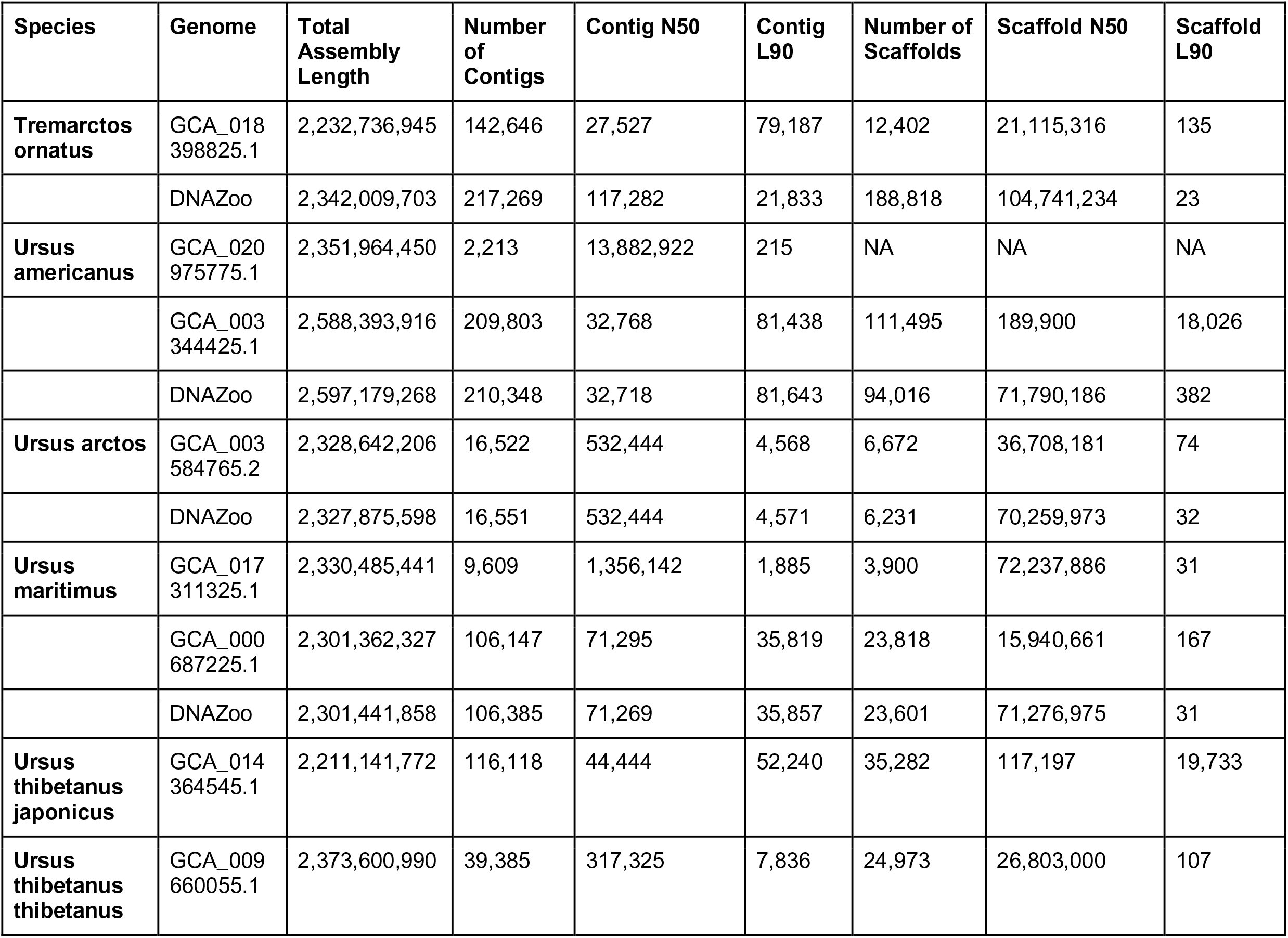
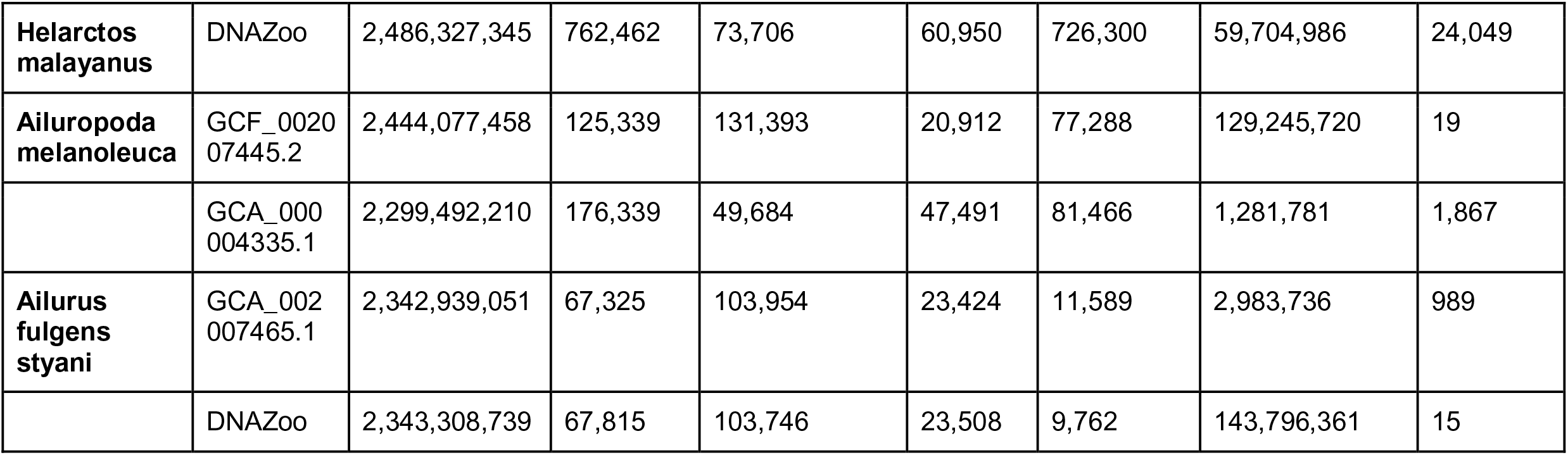
Assembly statistics for all other bear assemblies.

**Table S4:** BUSCO scores for intermediate and final brown bear assembly produced in this study using the mammalia_odb10 dataset.

| **Assembly** | **Complete** | **Single Copy** | **Duplicated** | **Fragmented** | **Missing** |
| --- | --- | --- | --- | --- | --- |
| **Hifiasm-Hap1** | 96.3% | 95.5% | 0.8% | 1.0% | 2.7% |
| **Hifiasm-Hap2** | 96.3% | 95.5% | 0.8% | 1.1% | 2.6% |
| **Hifiasm-Hap1 + HiC** | 96.5% | 95.7% | 0.8% | 1.0% | 2.5% |
| **Hifiasm-Hap2 + HiC** | 96.3% | 95.6% | 0.7% | 1.1% | 2.6% |
| **Composite** | 96.3% | 95.6% | 0.7% | 1.1% | 2.6% |

**Table S5:** BUSCO scores for all other bear assemblies (except for *Ursus arctos*, only assembly with highest continuity statistics, i.e. scaffold L90/N50, was analyzed).

| Species | Genome | Complete | Single Copy | Duplicated | Fragmented | Missing |
| --- | --- | --- | --- | --- | --- | --- |
| *Tremarctos ornatus* | DNAZoo | 94.9% | 94.1% | 0.8% | 1.6% | 3.5% |
| *Ursus americanus* | GCA_020975775.1 | 96.0% | 95.4% | 0.6% | 1.2% | 2.8% |
| *Ursus arctos* | GCA_003584765.2 | 96.1% | 94.6% | 1.5% | 1.2% | 2.7% |
|  | DNAZoo | 96.1% | 94.6% | 1.5% | 1.1% | 2.8% |
| *Ursus maritimus* | GCA_017311325.1 | 96.0% | 95.4% | 0.6% | 1.1% | 2.9% |
| *Ursus thibetanus japonicus* | GCA_014364545.1 | 80.1% | 79.6% | 0.5% | 8.4% | 11.5% |
| *Ursus thibetanus thibetanus* | GCA_009660055.1 | 95.9% | 94.3% | 1.6% | 1.1% | 3.0% |
| *Helarctos malayanus* | DNAZoo | 93.0% | 92.4% | 0.6% | 2.4% | 4.6% |
| *Ailuropoda melanoleuca* | GCF_002007445.2 | 94.8% | 92.8% | 2.0% | 1.5% | 3.7% |
| *Ailurus fulgens styani* | DNAZoo | 94.6% | 93.3% | 1.3% | 1.6% | 3.8% |

**Table S6:**
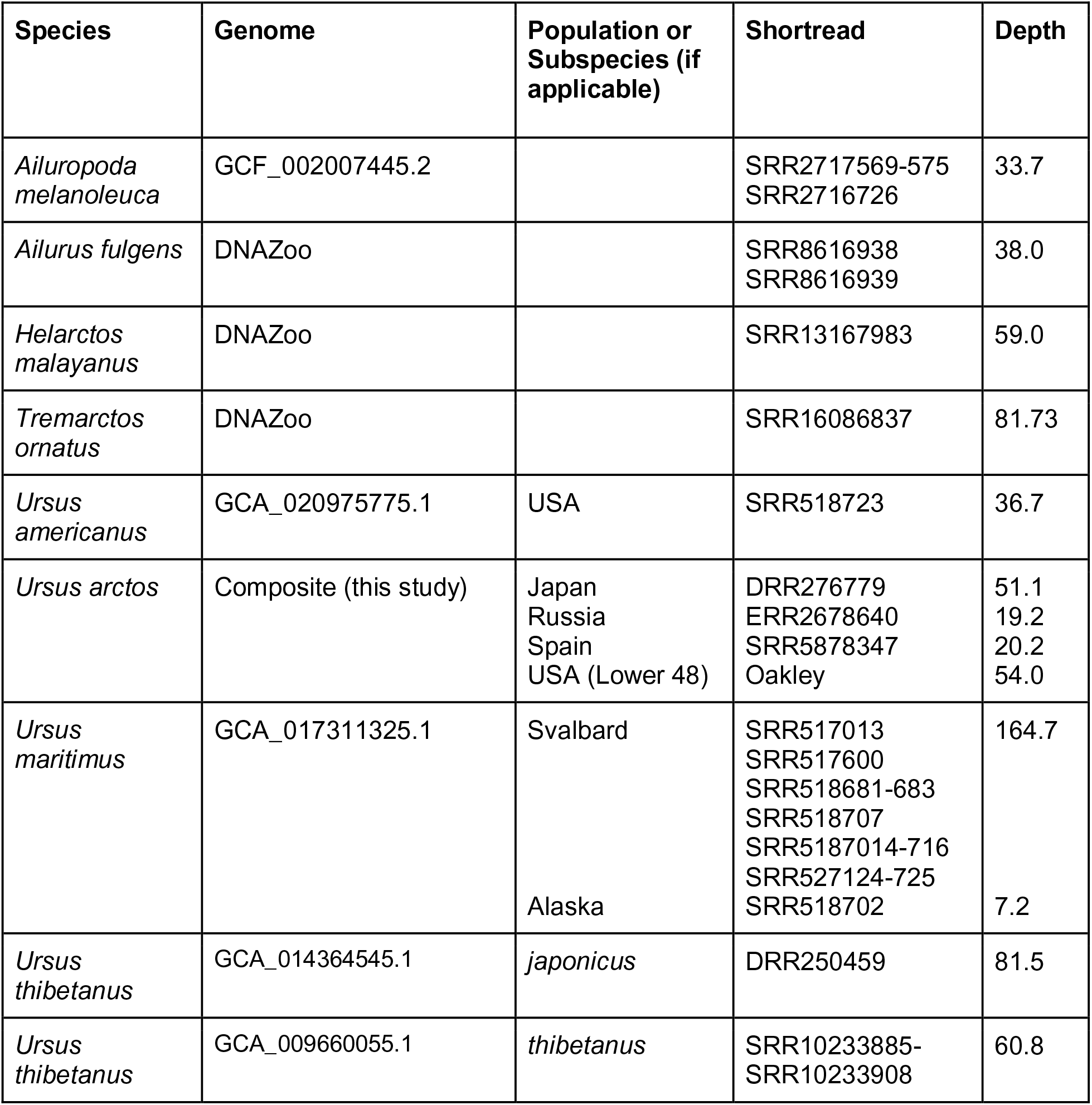
Data used in PSMC and heterozygosity analysis

